# Cold-induced loss of interaction with HSC70 triggers inflammasome activity of familial cold autoinflammatory syndrome-causing mutants of NLRP3

**DOI:** 10.1101/2022.11.16.516705

**Authors:** Akhouri Kishore Raghawan, Rajashree Ramaswamy, Ghanshyam Swarup

**Author notes:** **Corresponding authors:** Ghanshyam Swarup, Akhouri Kishore Raghawan. Akhouri Kishore Raghawan, Beth Israel Deaconess Medical Centre, Harvard Medical School, Boston, MA, 02115, USA.

## Abstract

NLRP3 is a cytoplasmic receptor protein, which initiates caspase-1 mediated inflammatory immune response upon detection of invading pathogen or a wide array of internal distress signals. Several gain-of function mutations of NLRP3 cause hereditary disorder of cold-induced hyper-inflammation known as familial cold autoinflammatory syndrome-1 (FCAS1). Although, caspase-1 activation and downstream interleukin-1β / interleukin-18 maturation are common effectors in pathophysiology of this disorder, molecular mechanisms of how exposure to subnormal temperature triggers mutant NLRP3-inflammsome activity is not understood. Here, we show that endogenous NLRP3 is in complex with HSC70 (HSPA8), and this interaction is reduced upon exposure to cold. FCAS-causing NLRP3-L353P and NLRP3-R260W mutants show enhanced interaction with HSC70. Upon exposure to subnormal temperature, NLRP3-L353P and NLRP3-R260W show enhanced inflammasome formation, increased caspase-1 activation and reduced interaction with HSC70. Knockdown of HSC70 results in increased inflammasome formation by L353P and R260W mutants of NLRP3. Our results suggest that interaction with HSC70 suppresses inflammasome formation by FCAS-causing NLRP3 mutants at physiological temperature, and loss of this inhibitory association at subnormal temperature causes aggravated inflammasome formation and caspase-1 activation leading to interleukin-1β maturation. These results provide evidence for HSC70 being a cold-sensor and a temperature-dependent regulator of inflammatory signaling by FCAS-causing NLRP3 mutants.

## 1. Introduction

NLRP3 (also known as cryopyrin, NALP3, CIAS1, PYPAF1) is a cytoplasmic pattern recognition receptor protein involved in innate immunity. NLRP3 induces inflammatory response upon detection of an invading pathogen, environmental stress or internal damage signals. In presence of an activating signal, NLRP3 initiates assembly of inflammasome, a multiprotein complex consisting of adaptor protein ASC, procaspase-1 and NEK7 [1-3]. Inflammasome acts as a platform for proximity-induced autoactivation of caspase-1, augmenting cleavage and maturation of inflammatory cytokines like IL-1β and IL-18. Gain-of-function mutations in NLRP3 lead to a constitutively active NLRP3 that induces unchecked inflammasome assembly and caspase-1 activation resulting in sterile inflammatory disease conditions referred to as cryopyrin-associated periodic syndromes (CAPS) [4]. CAPS is a disease continuum of heritable disorders of autoinflammatory phenotypes of periodic fever, general malaise, urticaria, renal amyloidosis and bilateral sensorineural hearing loss. Interestingly, different mutations in the NLRP3 affect a distinct set of phenotypes; moreover, different individuals carrying the same NLRP3 mutation may be phenotypically different. Familial cold autoinflammatory syndrome 1 (FCAS1) is a subset of CAPS, presented with temperature-dependent phenotype wherein the symptoms aggravate only when the patient is exposed to subnormal temperature [4-12]. Monocytes obtained from FCAS1 patients carrying NLRP3-L353P mutation show enhanced cytokine IL-1β release upon exposure to subnormal temperature, which is caspase-1 dependent [13-15]. Blocking of IL-1β itself or its receptor leads to reduction in symptoms in FCAS patients carrying NLRP3 mutations [16-22]. These and several other observations suggest that caspase-1 activation and consequent cytokine maturation and release play a crucial role in FCAS1 pathophysiology. However, the factors that suppress activity of FCAS1-associated NLRP3 mutants at physiological temperature but allow resumption of its activity at sub physiological temperature are not known.

Heat shock protein (HSP) family member chaperones HSP70 and heat shock cognate protein 70 (HSC70/HSPA8) have been linked to regulation of inflammatory signaling downstream of immune receptors NLRP3 and NLRC4. HSP70 has been shown to regulate inflammasome activity of wild-type NLRP3 while HSC70 negatively regulates activity of FCAS-associated mutant NLRC4-H443P [23,24]. HSC70 undergoes a temperature-dependent conformational change which leads to increase in its binding to client proteins with increase in temperature in the physiological temperature range of 30°C to 37°C [25]. These properties of HSC70 indicate that it is a potential candidate for temperature-dependent regulation of FCAS mutants. In this study, we have investigated the role of HSC70 in temperature-dependent regulation of inflammasome activity of two FCAS1-associated mutants, NLRP3-R260W and NLRP3-L353P.

## 2. Materials and Methods

### 2.1. Cell Culture, treatments and transfections

HEK293T cells were maintained in Dulbecco’s modified Eagle Medium (DMEM) containing 10% fetal bovine serum (FBS) while THP1 cells were maintained in RPMI supplemented with 10% heat inactivated FBS, in an incubator that maintained 37°C and 5% CO_2_. For exposure to subnormal temperature, cells were shifted to incubators maintained at 28°C for indicated duration. THP1 cells were differentiated into macrophage-like cells by treatment with 10nM phorbol 12-myristate 13-acetate (PMA) for a period of 72h. Lipofectamine2000 (cat no. #11668019) from Thermo Fisher Scientific or Helix-IN reagent (cat no. HX11000) from OZ Biosciences was used to transfect HEK293T cells as per the manufacturer’s protocol. In general, about 50% - 60% HEK293T expressing cells were observed in our experiments.

### 2.2. Antibodies and Chemicals

NLRP3 antibody (CST#13158) was from Cell Signaling Technology. GAPDH antibodies (MAB-374) and Actin (MAB1501), and Cy3 (Indocarbocyanine) conjugated mouse or rabbit secondary antibodies were purchased from Millipore. N-ethylmaleimide (cat no. E3876) and PMA (P1585) were from Sigma-Aldrich. Antibodies against caspase-1 (SC-56036), myc (SC-40; SC-789; SC-515), GFP (cat no. SC-9996), HSC70 (SC-7298). Haemagglutinin (HA; SC-805), IL-1β (SC-7884) and Blotto (sc-2324) were obtained from Santa Cruz Biotechnology, USA. HSC70-siRNAs (sc-29349) and control siRNA (sc-37007) were from Santa Cruz Biotechnology, USA. HRP-conjugated mouse (NA931) and rabbit (NA934) secondary antibodies were from GE Healthcare; FemtoLUCENT™ PLUS HRP Kit (cat no. 786-10) and protease inhibitor cocktail (cat no. 786-437) were procured from GBiosciences. Anti-mouse Cy3 conjugated secondary antibody (cat no. AP124C) and anti-rabbit Cy3 conjugated secondary antibody (cat no. AP132C) were procured from Merck. Anti-rabbit Alexa Fluor 488-conjugated secondary antibody (cat. No. A11008) and Anti-mouse Alexa Fluor 488 conjugated secondary antibody (cat no. A10680) were obtained from Thermo Fisher Scientific.

### 2.3. Expression vectors

NLRP3 expression vectors pEGFP-C2-NLRP3 (#73955) and pCDNA-Myc-NLRP3-R260W (#73956) were obtained from Addgene, USA. The plasmids pEGFP-C2-NLRP3-R260W and pEGFP-C2-L353P were generated by inducing CGA to TGG, and CTG to CCG substitutions in pEGFP-C2-NLRP3 expression construct using site directed mutagenesis. Expression vector pCDNA-Myc-NLRP3-WT was generated by inducing TGG to CGG substitution in pCDNA-Myc-NLRP3-R260W at desired site. Similarly, the construct pCDNA-Myc-NLRP3-L353P was generated by inducing CTG to CCG substitution in pCDNA-Myc-NLRP3-WT construct by site-directed mutagenesis. The nucleotide sequence of the constructs was confirmed by automated DNA sequencing.

### 2.4. Indirect immunofluorescence, microscopy and quantitation of inflammasomes

Myc-tagged constructs expressing wild type NLRP3 or its mutants were co-expressed with HA-tagged ASC in HEK293T cells plated on coverslips. For exposure to subnormal temperature, cells were shifted to incubator maintained at 28°C for 4h after 12h of transfection. After 16h of transient expression, cells were fixed and stained with HA- and Myc-antibodies. For HSC70 knockdown, transfections with 100 pmols of control siRNA or HSC70-specific siRNA was carried out in HEK293T cells plated on coverslips. After 24h of transfection, a second set of transfections was carried out with siRNAs and desired expression constructs of NLRP3 and ASC. Cells were fixed with formaldehyde after 42h of first transfection and stained for Myc- and HA-tags using specific antibodies. Immunostained cells were observed under fluorescence microscope to score for presence of specks in co-expressing cells [24]. Data are presented as mean±S.D. of % cells forming specks from at least 3 independent experiments performed on duplicate coverslips after examining at least 200 co-expressing cells from each coverslip. An automated AxioImager Z.2 (Zeiss) fluorescence microscope with Axiovision software and x40/0.75 NA dry objective was used to capture images.

### 2.5. Co-immunoprecipitation and western blotting

Immunoprecipitations and western blotting were carried out essentially as described [24]. Macrophage-like differentiated THP1 cells were lysed for 30minutes at 4°C, in buffer containing 150mM NaCl, 20mM Tris-HCl (pH7.5), 1mM PMSF, protease inhibitor cocktail, 0.5% NP40, 0.5mM EDTA, 0.1% BSA, and 10mM N-ethylmaleimide. Debris was removed by centrifugation (10000g for10minutes at 4ºC). Lysates were incubated with 2μg of NLRP3 antibody or control IgG (incubated with agarose conjugated Protein A/G for 2h at 4°C) 8h at 4°C on a Roto-torque. The immunoprecipitates were washed with buffer containing 150mM NaCl, 20mM Tris-HCl (pH7.5), 1mM PMSF, 0.5mM EDTA, protease inhibitor cocktail and 10mM NEM. After three washes, the immunoprecipitates were boiled in SDS containing Laemmli sample buffer for 5minutes. The samples were then western blotted to analyze bound proteins. For immunoprecipitation of GFP-tagged proteins, a GST-tagged GFP-nanobody purified from *E. Coli* BL-21 DE-3 cells was used and experiments performed as described earlier [24,26].

### 2.6. Quantification of western blots

Using ImageJ software (NIH), quantification of co-immunoprecipitated proteins was obtained by normalizing the intensities of bands corresponding to co-immunoprecipitated proteins with bands of immunoprecipitated proteins. For quantification of IL-1β maturation, intensities of mature IL-1β (p17) bands were normalized with pro-IL-1β signal (p32) as described [24].

### 2.7. Statistical analysis

Bar diagrams present quantitative data as mean±S.D. values. To determine significance of relative difference between a test sample and control (taken as 1), one-tailed Student’s t-Test was used, and two-tailed t-Test was used to determine significance of differences between two means.

## 3. Results

### 3.1. HSC70 complexes with cellular NLRP3 and shows enhanced interaction with FCAS-associated NLRP3 mutants

HSC70 is a chaperone protein involved in folding of large nascent polypeptides and refolding of misfolded polypeptides [27]. FCAS1-associated gain-of-function mutations in NLRP3 may cause alterations in the folding pattern of mutant NLRP3 polypeptides and this may alter interaction of the mutants with HSC70 (Fig.1*A*). Therefore, we examined the possibility of a molecular complex formation between HSC70 and NLRP3 and its FCAS1-associated mutants. The interaction of endogenous NLRP3 with HSC70 was examined in THP1 cells differentiated into macrophage-like cells by treatment with 10nM PMA for 72h. Western blot analysis of immunoprecipitates of cellular NLRP3 from lysates of differentiated THP1 cells showed that HSC70 forms a molecular complex with endogenous NLRP3 (Fig.1*B*). We further analyzed the interaction of NLRP3 and its mutants with endogenous HSC70. We carried out western blot analysis of immunoprecipitates of GFP-tagged wild type NLRP3 or NLRP3-R263W or NLRP3-L353P mutants obtained from lysates of HEK293T cells transiently expressing these proteins. We observed that compared to wild type NLRP3, FCAS-associated mutants of NLRP3, R263W and L353P, show significantly enhanced interaction with HSC70 (Fig.1*C* and *D*).

**Fig. 1.**
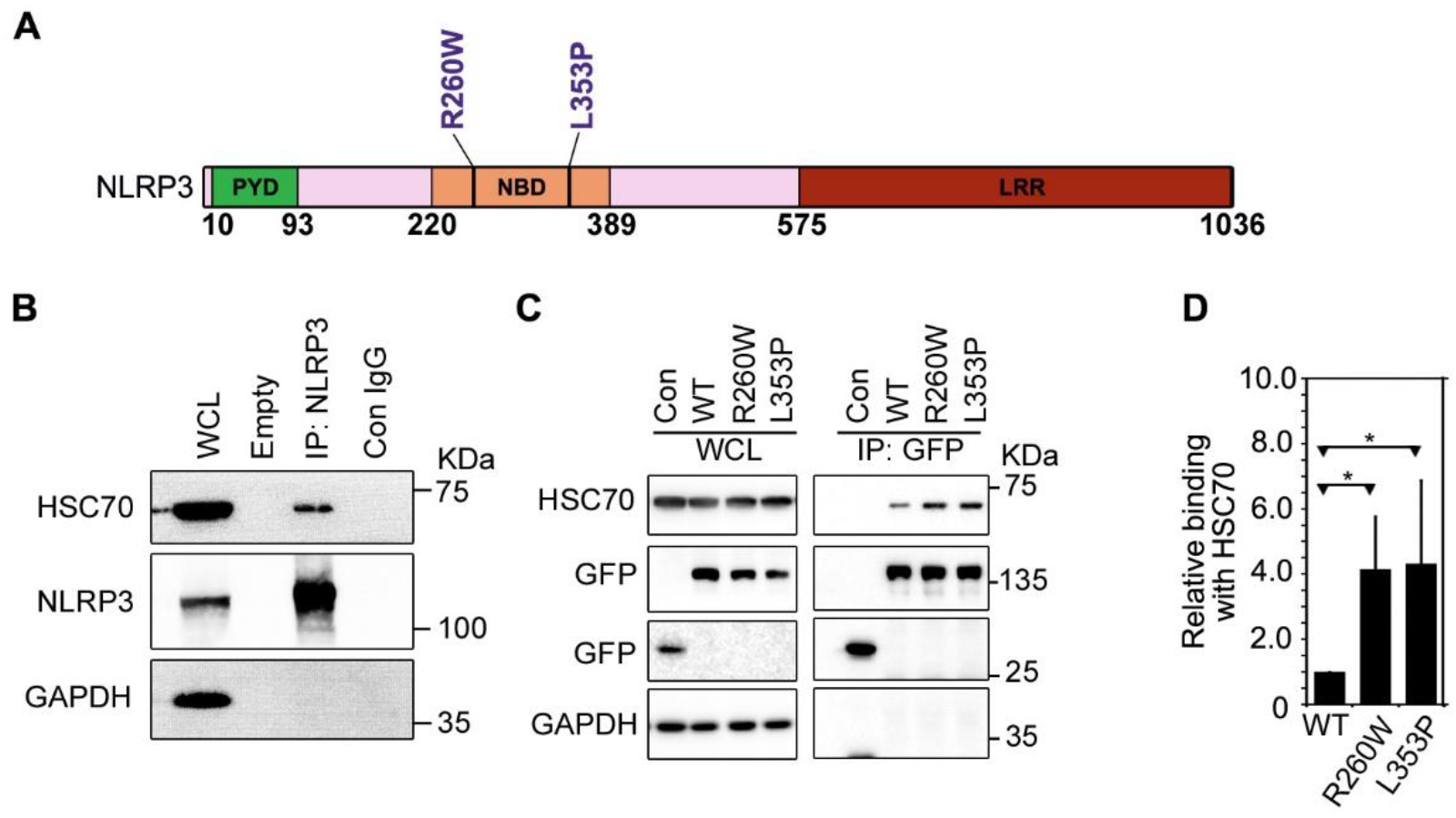
HSC70 complexes with cellular NLRP3 and shows enhanced interaction with FCAS-associated NLRP3 mutants. **(A)** Schematic showing domain organization of NLRP3. Sites of FCAS1-associated mutations in NLRP3 are indicated. PYD, pyrin domain; NBD, nucleotide-binding domain; LRR, leucine-rich repeat domain. **(B)** Endogenous NLRP3 forms a complex with HSC70. Western blot analysis of immunoprecipitates of NLRP3 obtained from lysates of differentiated THP1 cells shows interaction between NLRP3 and HSC70. Lysate was subjected to immunoprecipitation using NLRP3 antibody (IP: NLRP3) or normal IgG (Con IgG). WCL, whole cell lysate. GAPDH was used as a control to check for any non-specific binding. **(C)** Lysates of HEK293T cells transiently expressing GFP-tagged NLRP3 and its mutants were subjected to immunoprecipitation using GFP antibody. Western blot analysis showed presence of HSC70 in the immunoprecipitates (IP) of WT-NLRP3 as well as R260W and L353P mutants. **(D)** Bar diagram shows quantitation of relative abundance of HSC70 in IP of R260W and L353P mutant compared to WT-NLRP3 normalized with GFP signal from 5 independent experiments; n=5. * p<0.05.

### 3.2 HSC70 suppresses inflammasome formation by NLRP3 mutants at physiological temperature

An activated NLRP3 or its constitutively active mutants oligomerize with the adaptor protein ASC to form speck-like structures called inflammasomes. The intracellular ASC-speck formation is a marker of inflammasome activity of NLRP3 [28,29]. To test the role of HSC70 in regulating inflammatory signaling by NLRP3, we used specific siRNAs to knock down HSC70. The efficacy of siRNA to knockdown HSC70 in HEK293T cells was assessed by western blot (Fig. 2A). We co-expressed HA-tagged ASC with Myc-tagged wild type NLRP3, NLRP3-R260W or NLRP3-L353P in HEK293T cells plated on coverslips at 37°C. Compared to wild-type NLRP3, expression of NLRP3-R260W or NLRP3-L353P with ASC resulted in higher percentage of cells showing ASC-speck formation (Fig. 2B, C). ASC mediated speck formation by NLRP3-R260W as well as NLRP3-L353P further increased significantly upon HSC70 knockdown (Fig. 2B, C). Wild type NLRP3 mediated ASC-speck formation was not affected significantly upon knockdown of HSC70 (Fig. 2B, C). These results suggest that HSC70 is a suppressor of inflammasome formation by FCAS1-associated NLRP3 mutants R263W and L353P at physiological temperature.

**Fig. 2.**
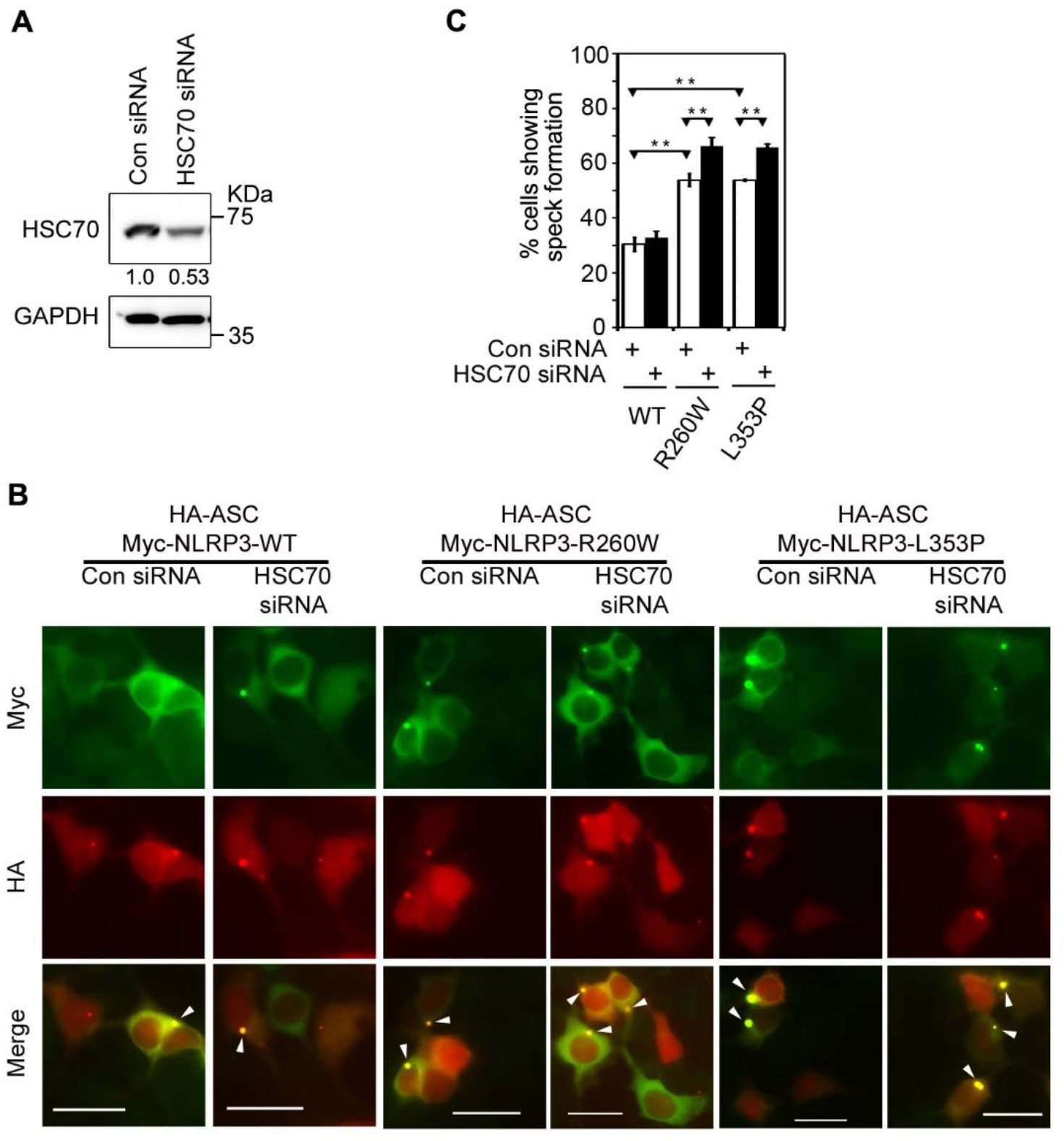
HSC70 negatively regulates inflammasome formation by NLRP3 mutants. **(A)** HEK293T cells were transfected with HSC70-specific siRNA or control siRNA (Con siRNA), and after 42h of transfection, siRNA mediated knockdown of HSC70 was assessed by western blotting. GAPDH was used as loading control. **(B)** Effect of HSC70 knockdown on speck formation by WT-NLRP3, NLRP3-R260W and NLRP3-L353P. Representative images show enhanced ASC-speck formation by NLRP3 mutants upon siRNA mediated HSC70 knockdown. Cells co-expressing HA-ASC and Myc-NLRP3, Myc-NLRP3-R260W or Myc-NLRP3-L353P were scored for presence of specks. ASC-specks formed inside the cells are indicated by white arrowheads. Scale bars, 10μm. **(C)** Bar diagram shows quantitation of effect of HSC70 knockdown on ASC-speck formation by NLRP3, NLRP3-R260W and NLRP3-L353P. n=3. ** p<0.005.

### 3.3. Effect of subnormal temperature on interaction of NLRP3 and its FCAS-associated mutants with HSC70

HSC70 undergoes a reversible conformational change upon exposure to subnormal temperature, thereby losing its ability to engage with its client proteins [25]. We tested the effect of exposure to subnormal temperature on the interaction of HSC70 with wild type NLRP3 and its FCAS-associated mutants. HEK293T cells expressing GFP-tagged wild type NLRP3, NLRP3-R260W or NLRP3-L353P were grown at 37°C for 16h or shifted to 28°C for 4h before lysis, and cell lysates were subjected to immunoprecipitation. These immunoprecipitates were analyzed by western blotting. Quantification of western blots showed significantly reduced interaction between HSC70 and wild type NLRP3, NLRP3-R260W or NLRP3-L353P upon exposure of cells to subnormal temperature (Fig. 3A, B). We also analyzed the effect of subnormal temperature on the interaction of endogenous NLRP3 with HSC70. THP1 cells were differentiated into macrophage cells using 10nM PMA for 72h. For exposure to subnormal temperature, one set of cells was incubated at 28°C for 6h after 66h hours of differentiation, and another set of cells was maintained at 37°C. Cell lysates were then subjected to immunoprecipitation using NLRP3 antibody. Western blot analysis of immunoprecipitates showed significant reduction in interaction between cellular HSC70 and NLRP3 at 28°C (Fig. 3C, D).

**Fig. 3.**
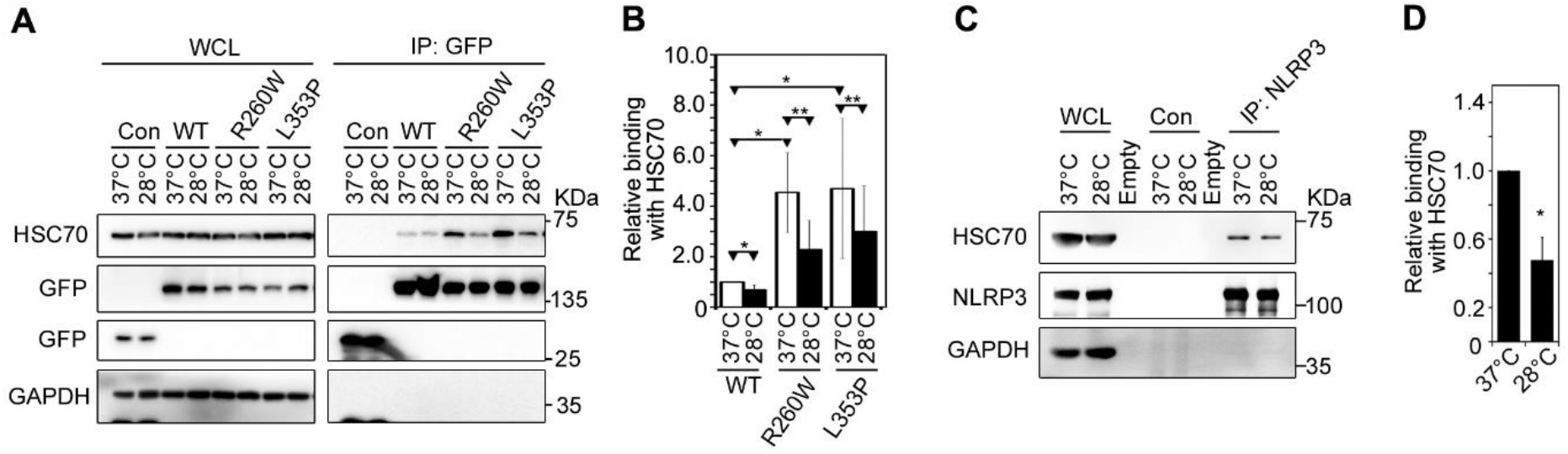
Effect of subnormal temperature on interaction of NLRP3 and its FCAS-associated mutants with HSC70. **(A)** HEK293T cells expressing GFP constructs of indicated plasmids were grown at 37°C for 16h or exposed to 28°C for 4h after 12h of transfection. Cell lysates were subjected to immunoprecipitation using GFP antibody and immunoprecipitates analyzed by western blotting. WT, wild-type NLRP3. **(B)** Quantitation of co-precipitated HSC70 at 37°C and at 28°C is shown. Values are normalized with corresponding IP signals. n=4. * p<0.05; ** p<0.005. **(C)** THP1 cells were treated with 10nM PMA for 72h at 37°C, or exposed to 28°C for 6h after 66h of treatment with PMA. Cell lysates were subjected to immunoprecipitation using NLRP3 antibody (IP) or normal IgG (Con) and immunoprecipitates analyzed by western blotting. **(D)** Bar diagram shows quantitation of relative binding of HSC70 with NLRP3 at 37°C and 28°C. n=3. * p<0.05.

### 3.4 Effect of exposure to subnormal temperature on inflammasome formation and caspase-1 activation by NLRP3 mutants

We further tested the effect of subnormal temperature on the ability of NLRP3 mutants to form ASC-specks. HEK293T cells plated on coverslips, and transiently expressing HA-tagged ASC along with Myc-tagged wild type NLRP3 or its mutants were incubated at 28°C for 4h after 12 hours of expression at 37°C, or left at 37°C for 16h. These cells were then scored for ASC-specks. NLRP3-L353P or NLRP3-R260W expressing cells showed increased ASC-speck formation compared to wild type NLRP3, which further increased significantly upon exposure of the cells to 28°C for 4h (Fig. 4A, B).

**Fig. 4.**
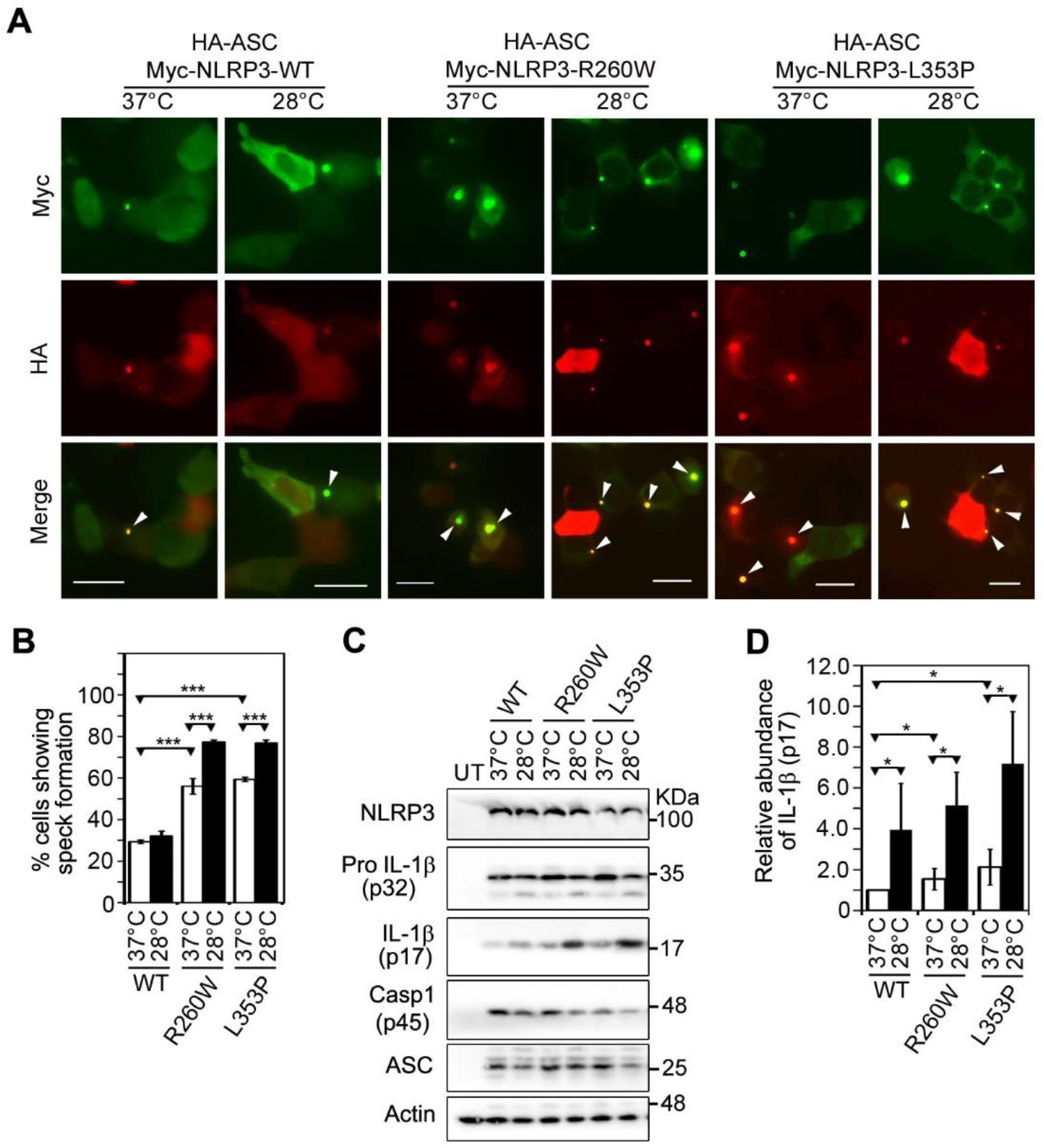
Effect of subnormal temperature on inflammasome formation and caspase-1 activation by NLRP3 mutants. **(A)** HEK293T cells coexpressing HA-ASC along with wildtype NLRP3, NLRP3-R260W or NLRP-L353P were scored for presence of ASC-specks. For cold-exposure, one set of cells was shifted to 28°C for 4h after 12h of transfection while the other set was maintained at 37°C. Representative immunofluorescence images show increase in percentage of cells forming ASC-specks upon exposure to subnormal temperature. White arrowheads indicate specks. Scale bars, 10μm. **(B)** Bar diagram shows quantitation of ASC-speck formation upon exposure of cells to subnormal temperature. n=6. *** p<0.0005. **(C)** Western blot analysis shows enhanced IL-1β maturation by NLRP3-R260W and NLRP-L353P mutants upon exposure of cells to 28°C. HEK293T cells were transfected with indicated NLRP3 plasmids along with caspase-1, ASC and pro-IL-1β. The transfected cells were grown at 37°C for 24h or incubated at 28°C for 6h after 18h of transfection. **(D)** Bar diagram shows quantitation of relative abundance to mature IL-1β (p17) normalized with the levels of pro-IL-1β (p32). n=5. * p<0.05.

We examined NLRP3 and its mutants for their ability to affect IL-1β maturation upon exposure of cells to subnormal temperature. IL-1β maturation is often used to estimate caspase-1 activation because pro-IL-1β is a very specific substrate of caspase-1. HEK293T cells were transfected with indicated NLRP3 construct, caspase-1, ASC and pro-IL-1β for 24h, one set was exposed to 28°C for 6h after 18h of transfection, and another set was maintained at 37°C for 24h. Western blot analysis showed significantly enhanced IL-1β maturation by NLRP3-R260W and NLRP3-L353P in comparison with wild type NLRP3, which further increased significantly upon exposure of cells to 28°C (Fig. 4C. D). These results show that inflammasome formation and caspase-1 activation by NLRP3-R260W and NLRP3-L353P are significantly enhanced upon exposure of cells to subnormal temperature.

## 4. Discussion

NLRP3 forms inflammasomes upon activation by a wide range of danger signals that include PAMPs (pathogen-associated molecular patterns), physiological wastes like uric acid crystals, cholesterol crystals, internal stress signals like high ATP levels, altered pH and increased ROS levels [1,2]. Gain-of-function mutations in NLRP3 can cause uncontrolled activation of NLRP3 inflammasome resulting in constitutive caspase-1 activation and cytokine maturation leading to autoinflammatory disorders [4]. A subset of NLRP3-inflammasomopathies, wherein the inflammatory activity of certain NLRP3 mutants remains suppressed until exposure to subnormal temperature is called FCAS1. In this study, we have provided evidence that cold-triggered inflammasome activation downstream of FCAS1-associated NLRP3-R260W and NLRP3-L353P mutants is controlled by HSC70. A suppressive interaction with HSC70 keeps the activity of mutant NLRP3 in check at physiological temperature; upon exposure to subnormal temperature (28°C), loss of interaction with HSC70 allows mutant NLRP3 to constitutively assemble inflammasome and activate caspase-1 that leads to maturation of inflammatory cytokine IL-1β.

The chaperone HSC70 is actively involved in folding of nascent polypeptides and refolding of misfolded proteins. HSC70 recognizes small patches of hydrophobic sequences in the client proteins and synthetic peptides [25,27]. It is possible that certain mutations in NLRP3 augment conformational changes such that it gets recognized as a misfolded polypeptide by HSC70 chaperone. Engagement with HSC70 prevents the mutant NLRP3 from forming active inflammasome complexes, resulting in diminished inflammatory signaling downstream. However, upon exposure to sub physiological temperatures, HSC70 undergoes reversible conformational changes, and in the process, loses its ability to engage with its client proteins [25]. Thus, HSC70 acts as a temperature-dependent regulator of inflammasome function of FCAS1-associated NLRP3 mutants, L353P and R260W. This is shown schematically in Fig. 5. The most important features of this proposed model are: (a) enhanced interaction of HSC70 with FCAS-causing NLRP3 mutants, (b) negative regulation of inflammasome function of NLRP3 mutants by HSC70 at 37°C, and (c) low temperature-dependent reduction in interaction of HSC70 with NLRP3 mutants leading to their activation.

**Fig. 5.**
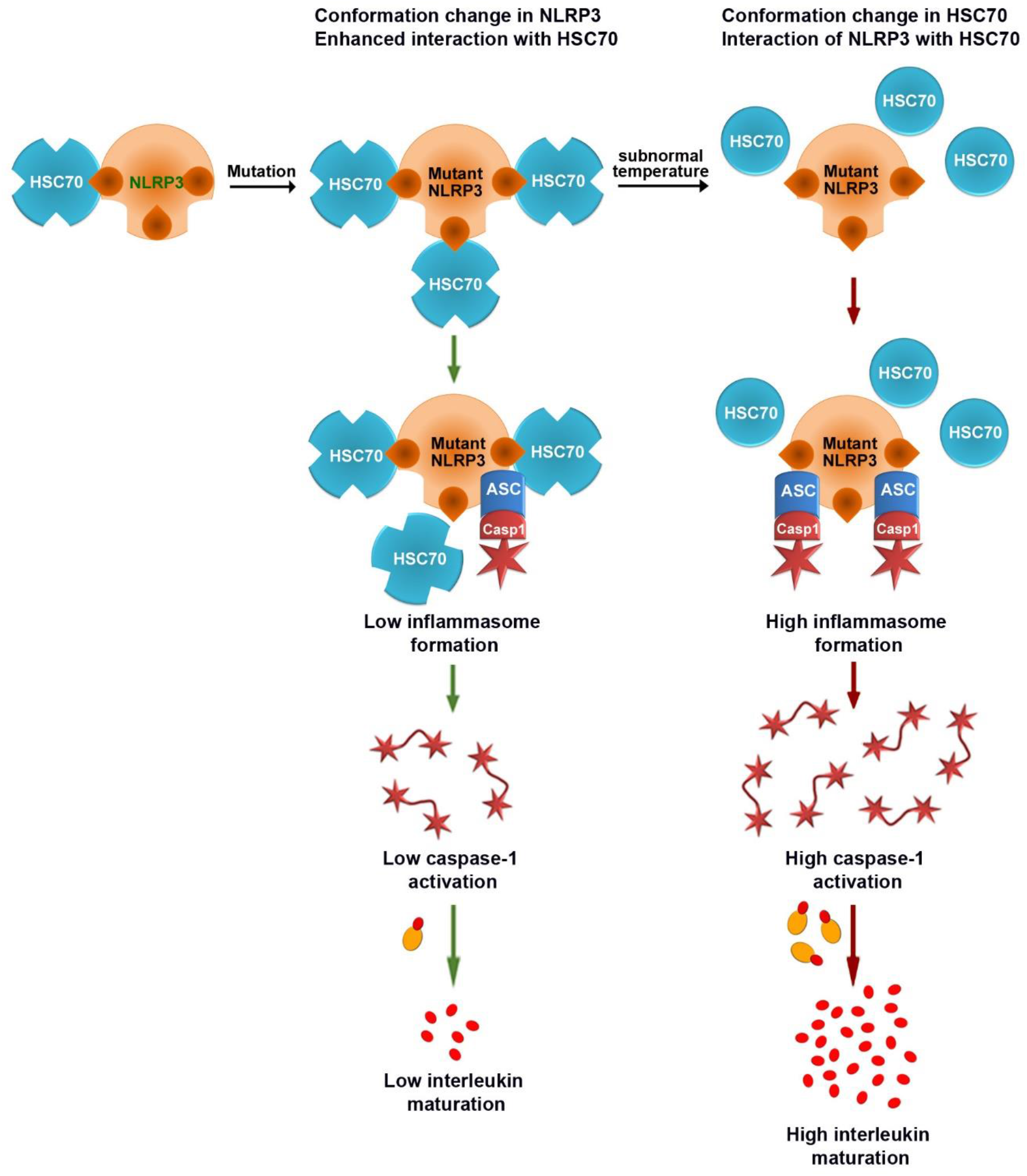
A model showing proposed mechanism of regulation of FCAS-causing mutants of NLRP3 by HSC70 in a temperature-dependent manner. Wild-type NLRP3 interacts with HSC70 and this interaction is increased by FCAS-causing mutations of NLRP3. It is proposed that FCAS-causing mutations induce a conformational change in NLRP3 that exposes HSC70-binding sites resulting in enhanced interaction of these mutants with HSC70. This interaction with HSC70 keeps the activity of NLRP3 mutants suppressed. Upon exposure to subnormal temperature HSC70 undergoes a conformational change resulting in loss of binding to the NLRP3 mutants. This leads to enhanced inflammasome formation and caspase-1 activation by the mutants resulting in enhanced cytokine maturation and release.

It is conceivable that in FCAS1, the mutant NLRP3 proteins themselves are cold sensors, which undergo temperature-dependent conformational change upon exposure to cold environment. In such a scenario every FCAS1 mutant must undergo a conformational change in the same temperature range (30-37°C), and this conformational change should lead to their activation.

However, such temperature-dependent conformational change has not been shown in any FCAS mutant so far. Our hypothesis proposes HSC70 as a common cold sensor, in which a temperature-dependent conformational change in the temperature range relevant for FCAS, is well documented [25]. In addition, temperature-dependent change in the interaction of HSC70 with client proteins is also well known [24,25,30]. Recently we shown that inflammasome activity of a FCAS4-associated mutant of NLRC4, H443P, is kept under control by HSC70 at 37°C, and upon exposure to low temperature, HSC70 dissociates from this mutant leading to inflammasome activation by this mutant [24].

The cold-sensitive ion channel TRPM8 could conceivably mediate activation of NLRP3 mutants through calcium ion mediated signaling. However, TRPM8 responds to a temperature range (8-26°C) which is different from that is involved in FCAS [31]. TRPM8 activation by chemicals leads to enhanced anti-inflammatory cytokine release [32]. Therefore, TRPM8 is not likely to be a cold sensor for FCAS1. Schade and colleagues have explored the mechanism of temperature dependent regulation of enzyme activity of FCAS3-causing PLCG2 mutants [33]. They identified the regions on PLCG2 molecule possibly involved in this temperature dependent regulation, and suggested that cold sensitive ion channels such as TRPM8 and TRPA1 are not likely to be involved in cold sensing in FCAS3 because these channels respond to much lower temperatures.

We have earlier provided evidence to show that HSC70 acts as a temperature sensor in the pathology of cold-triggered inflammatory signaling downstream of NLRC4-H443P mutant [24]. We have recently suggested that HSC70 could be a bonafide cold-sensor involved in the pathogenesis of FCAS disease spectrum, that involves mutations in NLRP3, NLRP12, PLCG2 and NLRC4 genes [34]. Through these results, we have added important evidence for HSC70 being a common cold-sensor and a temperature-dependent regulator of inflammatory signaling in FCAS. Our study highlights an important, but understudied property of HSC70 being a cold-sensor, and its role in regulating temperature-dependent signaling.

## Author contributions

AKR planned the experiments, performed the experiments, analyzed the data and wrote the paper. RR performed the experiments. GS conceived the study, planned the experiments, analyzed the data and wrote the paper. All authors approved final version of the manuscript.

## Acknowledgements

This work was supported by the J.C. Bose National Fellowship (grant no: SR/S2/JCB-41/2010) to GS from the Science and Engineering Research Board, Department of Science and Technology, Government of India, and Senior Scientist position to GS by the Indian National Science Academy, New Delhi.

## Competing interests

The authors declare that they have no competing interests.

## Notes

### Competing Interest Statement

The authors have declared no competing interest.

